# FlexAID: Revisiting docking on non native-complex structures

**DOI:** 10.1101/011791

**Authors:** F. Gaudreault, R. Najmanovich

## Abstract

Small-molecule protein docking is an essential tool in drug design and to understand molecular recognition. In the present work we introduce FlexAID, a small-molecule docking algorithm that accounts for target side-chain flexibility and utilizes a soft scoring function, i.e. one that is not highly dependent on specific geometric criteria, based on surface complementarity. The pairwise energy parameters were derived from a large dataset of true positive poses and negative decoys from the PDBbind dataset through an iterative process using Monte Carlo simulations. The prediction of binding poses is tested using the independent Astex dataset while performance in virtual screening is evaluated using a subset of the DUD dataset. We compare FlexAID to AutoDock Vina, FlexX, and rDock in an extensive number of scenarios to understand the strengths and limitations of the different programs as well as to reported results for Glide, GOLD and DOCK6 where applicable. The most relevant among these scenarios is that of docking on flexible non native-complex structures where as is the case in reality, the target conformation in the bound form is not known *a priori.* We demonstrate that FlexAID, unlike other programs, is robust against increasing structural variability. FlexAID obtains equivalent sampling success as GOLD and performs better than AutoDock Vina or FlexX in all scenarios against non native-complex structures. FlexAID is better than rDock when there is at least one critical side-chain movement required upon ligand binding. In virtual screening, FlexAID rescored results are comparable to those of AutoDock Vina and rDock. The higher accuracy in flexible targets where critical movements are required, intuitive PyMOL-integrated graphical user interface and free source code as well as pre-compiled executables for Windows, Linux and Mac OS make FlexAID a welcome addition to the arsenal of existing small-molecule protein docking methods.

**AUTHOR SUMMARY:** Protein ligand interactions are essential to understand biological processes such as enzymatic reactions, signalling pathways as well as in the development of new medicines. Docking algorithms permit to predict the structure of a ligand protein complex at the atomic level. Several docking algorithms were developed over the years with a tendency towards utilizing very specific and detailed (i.e., hard) descriptions of molecular interactions. In this work we present a new docking algorithm called FlexAID that utilizes a very general and superficial (i.e., soft) description of interactions based on atomic surface areas in contact. We demonstrate that FlexAID can achieve better accuracy in predicting the structure of ligand protein complexes than existing accessible widely used or state-of-the-art methods in real scenarios when using flexible targets harbouring structural differences with respect to the final protein structure present in the ligand protein complex. FlexAID and its PyMOL-integrated graphical user interface are free, easy to use and available for Windows, Linux and Mac OS.

## INTRODUCTION

Proteins co-exist as an ensemble of states in thermodynamic equilibrium [1]. The fraction of the population in each state is proportional to its free energy difference with respect to some reference state. Protein structure is subject to crystal packing constraints and may vary among different crystals of the same protein [2, 3]. At times such constraints may lead to distinct arrangements of side-chains in the binding-site [4]. The structure of a ligand-protein complex observed in a crystal represents the most populated state in the ensemble and often less-populated alternative states can be observed. Side-chain flexibility is common upon ligand binding [5] and in approximately 30% of cases, side-chain rotamer changes between Apo and Holo forms are essential for ligand binding [6]. According to the conformational selection theory for ligand-protein binding [7], a ligand that preferentially binds a given state will shift the equilibrium towards that state. Therefore, the bound structure may not necessarily represent the most common state of the protein when unbound due to a combination of the factors above.

Molecular docking is the most commonly used computational method to predict the structure of ligand-protein complexes. Many popular docking algorithms such as FlexX [8, 9], GOLD [10], DOCK6 [11] and AutoDock Vina [12] use stringent geometric constraints to define molecular interactions between the two molecules, particularly H-bonding interactions. Their scoring functions are called hard scoring functions [13] as good scoring depends on meeting stringent geometric constraints. These constraints assume that the protein side-chains are already correctly oriented in the binding-site to accommodate ligand binding. If these are not properly oriented, a finer search of the terminal atoms is thus required to satisfy the required geometric criteria. With some notable exceptions [14, 15], algorithms tend to be benchmarked using sets of ligand-bound (holo form) protein structures [16-18]. We call such complex structures native in the sense that the holo form conformation is the one that corresponds to the one observed with a bound ligand. Using this nomenclature, a non-native conformation is one obtained from an Apo (unbound) form of the protein or non-Apo form, i.e., a holo form bound to a ligand other than the one present in the native form being used as reference. Thus, most algorithms are tested using proteins in rather convenient conformations and do not discuss the accuracy of their method when tested on non native-complex structures. This scenario is not representative of a real-life situation considering that docking is mostly used to accelerate drug discovery or understand potential ligand-protein interactions where the protein structure considered often is a homology model or a non native-complex structure with respect to the ligand of interest. Additionally, protein flexibility is often neglected in virtual screening experiments, assuming that all ligands preferentially select the same state in the conformational ensemble [19].

Different levels of protein flexibility are observed when a ligand binds. Zavodsky *et al.* [20] suggested that side-chains move as minimally as possible to accommodate ligand binding, a concept they named the minimal rotation hypothesis. We recently quantified the minimal rotation hypothesis and observed that it is valid in approximately 20% of cases [6]. In about 90% of binding-sites, at least one side-chain rotamer change is observed in the bound complex [6]; and in 30% of these cases, side-chain movements are essential as severe steric clashes are observed in the holo form were it to retain the apo-form side-chain conformations. In the present study, we introduce the molecular docking algorithm FlexAID that uses a scoring function based on surface complementarity. The use of surface complementary in a docking scoring function was previously introduced in the LIGIN docking method [21, 22]. Rather than using stringent geometric criteria to define interactions, the method focuses on maximizing shape complementarity between atoms of the ligand and the protein representing favourable interactions as defined by a set of atom types and a pairwise atom type interaction matrix. We investigate to what extent a smooth scoring, i.e. one whose values do not change abruptly with slight changes in the structure, can implicitly simulate protein flexibility accommodating minimal side-chain rearrangements to take advantage of the higher success rates observed when soft potentials are used to mimic protein flexibility [11]. We also introduce explicit side-chain rotamer changes in our method to account for the larger movements that may lead to more drastic changes in the predicted energy and may not be covered by a smooth function.

The development of scoring functions is often based on the use of large datasets of complexes and ignores the potential contribution that negative examples may provide. For instance, empirical scoring functions such as the ones from AutoDock Vina, FlexX and rDock calibrate their energetic terms only using structures representing native complexes. Although the combined use of negative and positive examples was observed to be detrimental in other fields such as protein-protein interactions and protein structure predictions [23-25], it has been recently shown that it can improve the accuracy of scoring functions for protein-ligand docking [26]. The FlexAID method introduced here utilizes a soft scoring function, i.e., one that have no angular terms and does not vary abruptly with changes in geometry [13] developed using both positive and negative examples.

## RESULTS

Any optimization problem, docking included, has three interconnected components that affect the accuracy as well as usability of a method. These are representation, search and scoring.

FlexAID uses PDB structures ‘as is’, that is no manual curation is required. In particular, the introduction of hydrogen atoms prior to docking is not necessary and any such atoms are ignored if present. In addition, the calculation of partial charges or protonation is not required either. FlexAID relies on genetic algorithms as its search strategy as discussed in the methods section. The simplicity of the representation is directly interconnected with the scoring methodology and further details are given below.

### Development of the scoring function

We developed a soft scoring function for FlexAID using contact surfaces based on the complementarity function (CF) originally introduced by Sobolev *et al.* [22]. Unlike potential functions that describe the interaction between two atoms with a functional form with a minimum around an optimal distance, in the CF the interaction energy between two atoms varies linearly with their surface area in contact. As a soft scoring function, hydrogen atoms are implicitly accounted for. While this implicit treatment prevents the use of directionality constraints for interactions, the ‘softer’ or smoother surface helps maximize shape complementarity between protein ligand atoms. As described in the methods section, our implementation of the CF function is composed of three terms (Equation 1), a term to prevent steric clashes, a term representing interactions between ligand atoms and the implicit solvent and lastly, a term representing the interactions between ligand and protein atoms.

Ligand-protein interactions are quantified with a matrix of pairwise interaction energy terms between atom types and modulated by the surface area in contact between atoms. The pairwise parameters are derived with the use of statistical potentials optimized using a decoys set generated from the PDBbind database [27]. We grouped the proteins by sequence to account for the redundancy in the database so as not to bias the potentials towards pairwise atom type interactions frequently found in protein families more widely represented in PDBbind.

A Monte Carlo procedure was utilized to derive different potentials representing the 820 pairwise interactions between 40 atom types as described in the methods section. We iteratively made the decoy sets harder by injecting new low-energy decoys predicted with the potential from the preceding step. In order to make the decoys in each successive Monte Carlo iteration progressively more difficult, in addition to the increase of the number of decoys per complex, we also increased the number of complexes used to generate such decoys. With progressive iterations we increased the number of complexes that can be predicted while maintaining high area under the receiver-operator-curve (AUC) values on an independent test set (Table 1). The use of the PDBbind database to optimize the pairwise energy values makes the number of proteins considered in our study smaller than what was used in previous studies [26] but allows us to focus on a more relevant portion of the PDB in terms of drug-likeness of the ligands. We obtained an AUC around 0.6 when the parameters are set to zero (negative control), suggesting that the true positives have less steric clashes than the false positives as expected. Likewise, random values resulted in AUC values around 0.5. As expected, we observe that the AUC decreases slightly as the decoy sets are made larger and more difficult in successive MC iterations. Despite this slight decrease in AUC to a final value of 0.82, more complexes in the independent set can be successfully predicted (Figure 1). The obtained parameters also permit to better rank the solutions (Figure 1).

**Figure 1.**
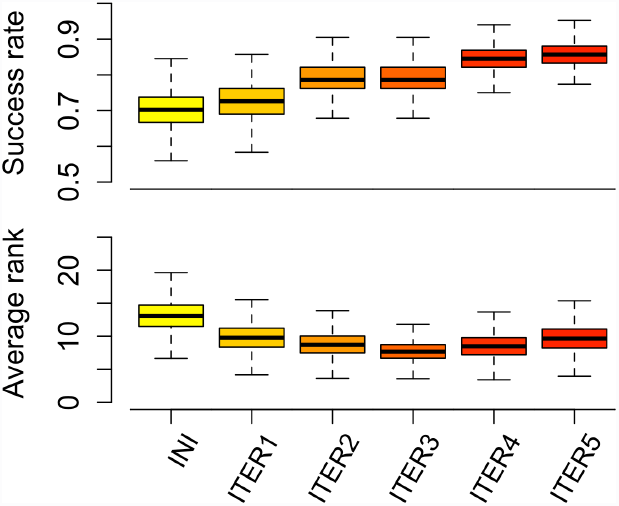
Bootstrapped average success rate and rank of FlexAID predictions on the independent Astex diverse set with consecutive parameter sets. The dataset comprises 84 protein-ligand complexes. The colours are used for clarity purposes only from the less optimized (yellow) to the more optimized (red) potential. The success rate is defined as the fraction of complexes where at least one successful solution (RMSD of 2.0Å from the native pose) was found. The rank is a measure of how well our method can discriminate the true binding mode from other modes. It represents the rank at which the first success is observed when the results are ordered according to the scoring function. The results are bootstrapped over 10 000 iterations.

**Table 1.** AUC for different parameter sets as a function of the number of Monte Carlo iterations on an independent dataset.

|  |  | Complexes | Groups | Decoys | AUC |  |  |
| --- | --- | --- | --- | --- | --- | --- | --- |
|  |  |  |  |  | Null<sup>a</sup> | Random<sup>b</sup> | Optimized<sup>c</sup> |
| MC Iterations | 1 | 1,207 | 544 | 310,459 | 0.586 | 0,526 ± 0,056 | 0.846 |
|  | 2 | 1,455 | 630 | 402,043 | 0.606 | 0,538 ± 0,046 | 0.842 |
|  | 3 | 1,543 | 657 | 455,047 | 0.611 | 0,514 ± 0,052 | 0.832 |
|  | 4 | 1,611 | 678 | 529,571 | 0.606 | 0,545 ± 0,061 | 0.820 |
|  | 5 | 1,649 | 688 | 592,385 | 0.604 | 0,529 ± 0,053 | 0.816 |

<sup>a</sup> All pairwise parameters set to 0.
<sup>b</sup> Average over a 10 random sets of parameters
<sup>c</sup> AUC values for the best matrix obtained in the given iteration

Given the functional form of the CF (Equation 1), the energy parameters obtained through the optimization process can be compared to our qualitative understanding of known chemical properties. To do so, we analysed the interactions between some of the most frequently occurring atom types in the training and testing datasets (Table S1). The top 14 pairwise interactions analysed account for 28% of the total contact surfaces observed in native structures. We assigned each interaction as either favourable or unfavourable based on chemical intuition (e.g., hydrophobic-polar interactions are unfavourable). We observe an ever-increasing agreement between the expected qualitative nature of interactions and their relative strength in the derived potential as the number of Monte Carlo iterations grows (Figures S1A-F). The final parameter set naturally encodes this chemical intuition (Figure 2). The interactions between positively and negatively charged atoms and other polar interactions are highly attractive while interactions between similarly charged atoms or pairs of donors or acceptors are found to be repulsive. In addition to this qualitative agreement, the relative ordering or the strength of interactions also makes sense from a chemical point of view. For example, the decreasing trend in the strength of polar interactions reflects an intuitive order in which interactions involving charged groups are stronger than weaker Hydrogen bonds. Moreover, the set of parameters naturally simulate the hydrophobic effect [28] as the exposure of polar atoms to solvent was observed to be less penalizing than the exposure of non-polar atoms. It is important to stress that this agreement between the strength of pairwise interaction energies and chemical intuition is a property of the interaction matrix that emerged as a natural result of the optimization process without being part of the optimization objective function.

**Figure 2.**
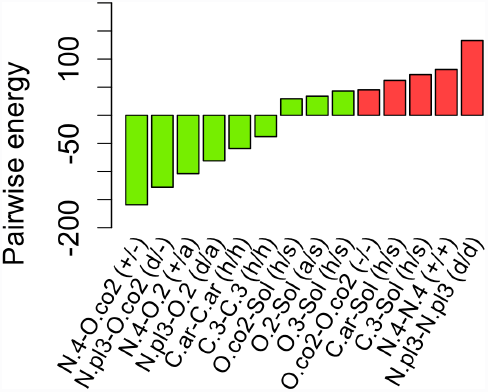
Pairwise interaction parameters for selected interactions. The lower a parameter, the more attractive is the interactions. Interactions that are attractive or repulsive according to chemical intuition are coloured in green or red respectively and shown in parenthesis: positive (+) or negative (-) charge, H-bond donor (d) or acceptor (a), hydrophobic (h) and hydrophilic (s). The following Sybyl atom types are shown: aromatic carbon (C.ar), aliphatic carbon (C.3) positively charged nitrogen (N.4), trigonal planar nitrogen (N.pl3), oxygen in carboxylate (O.co2), oxygen of carbonyl (O.2), oxygen of hydroxyl or ether (O.3) and solvent (Sol).

Interestingly, with the current approach the resulting strength of interactions between atom types and the solvent arises from the optimization process as repulsive independently of the atom type and validates previous approaches that added such solvent repulsive interactions ad-hoc [29]. In the context of molecular docking, repulsive solvent interactions force the ligand to interact with the target rather than maintain a large solvent accessible surface. It was previously found that optimizing for pose fidelity alone tend to increase the polar contributions of the scoring function [30]. Our results agree with this previous result as we also observe that non-polar interactions are generally weaker than polar interactions.

Docking algorithms estimate the energy of thousands of unique protein-ligand conformations during a simulation. Ideally, the scoring function would easily distinguish the native conformation from others and rank it as that with the lowest energy. We optimized the interactions such that the native conformation represents in as much as possible the global minimum of the energy landscape of the complementarity function. With the non-optimized interactions, we observe that the predicted energy of the best conformation (referred as *CF*_*min*_) suggested by our method is lower than the CF of the native conformation (*CF*_*ref*_) for nearly all complexes of the Astex diverse set [15, 31] that is independent from the PDBbind set (Figure S2A). This result show that the native conformations do not represent the global minimum of the CF indicating the presence of many local minima lower in energy than the native structure. This bias towards over-optimized conformations is drastically reduced as we iteratively improve the interaction matrix (Figures S2A-F), suggesting that the native conformations are increasingly the most favourable ones. This property again emerges naturally as a result of using positive and negative decoys and was not part of the optimization objective function. A few outliers with over-optimized CF are notable: PDB codes 1JD0, 1MZC, 1OQ5 and 2BR1. In 3 cases out of 4 cases, the discrepancies observed between *CF*_*min*_ and *CF*_*ref*_ can be explained by the presence of a metal coordinating the ligand, a type of interaction frequent in the Astex dataset but seldom observed in the PDBbind subset used for training (Table S1).

Complex conformations close to the native-complex structure (near-natives) share common and essential features of molecular recognition. Therefore, it is critical for the scoring function to also rank favourably these conformations to efficiently guide the search towards the true positive complexes. Conceptually, the energetic well must be smooth enough such that subtle changes in the structure do not drastically impact the overall energy. In order to do so, we assign decoys that are sufficiently close to the native complex structure in terms of RMSD as positive examples. We include at least 2 such designed true positives for each complex in the decoys set to build in smoothness into the parameter set. In Figure 3, we plot the relative difference in CF of all successful predictions in the Astex diverse set relative to the native CF. The relative CF difference is defined as Δ*CF*_*norm*_ = (*CF*_*min*_ − *CF*_*ref*_/*CF*_*ref*_. As a consequence of building the decoy sets in such a way, the resulting interaction matrix obtained after the successive Monte Carlo iterations displays a funnel-shaped distribution where larger deviations from the native conformation are associated with larger discrepancies in CF. This property of the CF function is notably less prominent in the non-optimized interaction matrix (Figure S3A). The outliers observed in the lower part could be partially explained by the RMSD measure itself. Some authors previously noted that the RMSD measure can be problematic as low RMSD poses may represent significantly different interactions in comparison to the native conformation [32, 33]. In fact, among the 90 complexes with Δ*CF*_*norm*_ < -0.25 (representing highly similar surfaces in contact), we find 80 complexes with RMSD > 1.5Å.

**Figure 3.**
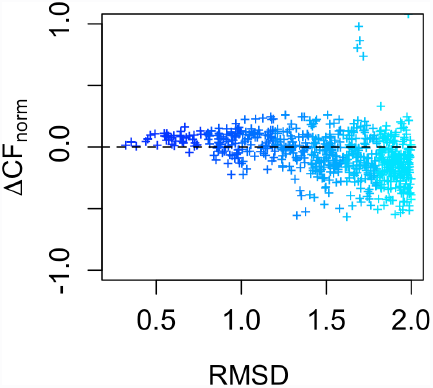

Smoothness of the FlexAID soft scoring function. As relative difference in CF is normalized by the CF of the reference native pose (*CF*_*ref*_ < *0*), complexes (represented by points) above the dashed line are cases where the CF of the best solution (*CF*_*min*_) is lower than that of the reference. The colours are used for clarity purposes only, from less (blue) to more distorted (cyan). The outliers observed above the dashed line represent cases where metals are present in the binding-site.

### Validation of FlexAID

We validate FlexAID extensively with respect to binding mode prediction as well as in virtual screening experiments and compare the performance of FlexAID to that of AutoDock Vina [12], FlexX [9] and rDock [18] and to reported results for DOCK6 [11], GOLD [15] and Glide [34].

#### Binding mode prediction

The performance of a docking algorithm is often evaluated by measuring how accurately the method can successfully predict, within a given margin of error, the native pose of the ligands for an ensemble of protein-ligand complexes. We compare our method to AutoDock Vina, FlexX and rDock for a number of reasons: These programs are freely available and widely used (AutoDock Vina and FlexX) or represent the state-of-the-art in docking simulations (rDock). All these programs utilize hard scoring functions. We compare the different software in various scenarios for complexes in the Astex diverse set [15, 31]. We use success rate to evaluate performance. The success rate measures the number of cases in a dataset where a successful pose is found within the top 10 scoring results. At times authors include all predicted poses in the definition of success rate (called sampling success) to report the performance of docking methods [18]. The use of sampling success as a measure of quality of a docking program is disingenuous, as it cannot be expected that users inspect all generated results. We believe reporting success rate among top 10 results is a reasonable compromise but we also calculated top 1, 5 and 100 success rates (Table S2). The later as an approximation of sampling success that would otherwise be measured over different numbers of solutions for the different methods. Some of the scenarios used to compare the different programs are not relevant in practical applications of docking simulations. Such scenarios highlight the factors that affect the performance of different docking programs tested and provide insight for their use in increasingly more real scenarios. Lastly, each method is tested on its own largest possible subset of the Astex diverse set for which the particular program could be run without technical errors (called here the ideal subset). Table S2 also shows the results obtained with the largest common subset of cases for all methods. The results are bootstrapped[35] over 10000 iterations to obtain a better approximation of average success rates across methods and do not differ significantly between the ideal and largest common subsets.

#### Rigid ligands on native structures

This experiment is not representative of a real-life scenario as neither the conformation of ligands nor that of the target are generally known prior to binding in a situation where docking is used to predict the structure of a protein-ligand complex. However, this experiment is useful in identifying cases that remain challenging even as the search space is reduced to optimizing the relative position of the protein and ligand as two rigid bodies. In this case, nearly all complexes (95.2%) can be successfully predicted with FlexAID (Figure S4A). In the same experiment, rDock attains a success rate of 93.2%. It is worth pointing out that the success rate obtained with rDock may be overestimated as the simulations were performed with the files provided by the authors for reproducing their results and do not include those for 11 proteins from the Astex diverse set that the authors chose not to provide. It is intriguing that AutoDock Vina predicts poorly (13.1%) when the ligand is treated as rigid despite docking into receptors that fit perfectly the ligand geometrically. The AutoDock Vina energy function uses strict criteria to describe atomic interactions with an angular term for H-bond interactions and strongly emphasizes directed interactions [13]. This result suggests that the hard scoring function in AutoDock Vina has a rough energy landscape and requires a very accurate search to find the specific geometric parameters required to achieve good scores. Similar to AutoDock Vina but to a lesser extent, the energy landscape of FlexX is also rough as it obtains a significantly lower success rate (71.8%) than FlexAID or rDock despite the restricted size of the search space.

#### Flexible ligands on native structures

In this case we again docked ligands into the native structure of proteins from the Astex diverse set but introduce ligand flexibility. Such a scenario is still somewhat artificial but would be representative of the approximately 10% of binding-sites that do not undergo side-chain conformational changes upon ligand binding [6]. The performance of FlexAID (67.9%) when docking in native structures is very similar to the one of AutoDock Vina (69.7%) (Figure 4A). FlexX and rDock achieve higher success rates of 78.8% and 87.8% respectively in this specific scenario where side-chain flexibility is not an issue. The sampling success of FlexAID is 85.7% compared to 93.4% for AutoDock Vina, 84.7% for FlexX and 98.6% for rDock. It is very surprising that it only takes the addition of ligand flexibility to so drastically improve the success rate of AutoDock Vina. This suggests the interplay between search and scoring helping to overcome the difficulties of finding good scoring solutions in rough energetic landscapes.

**Figure 4.**
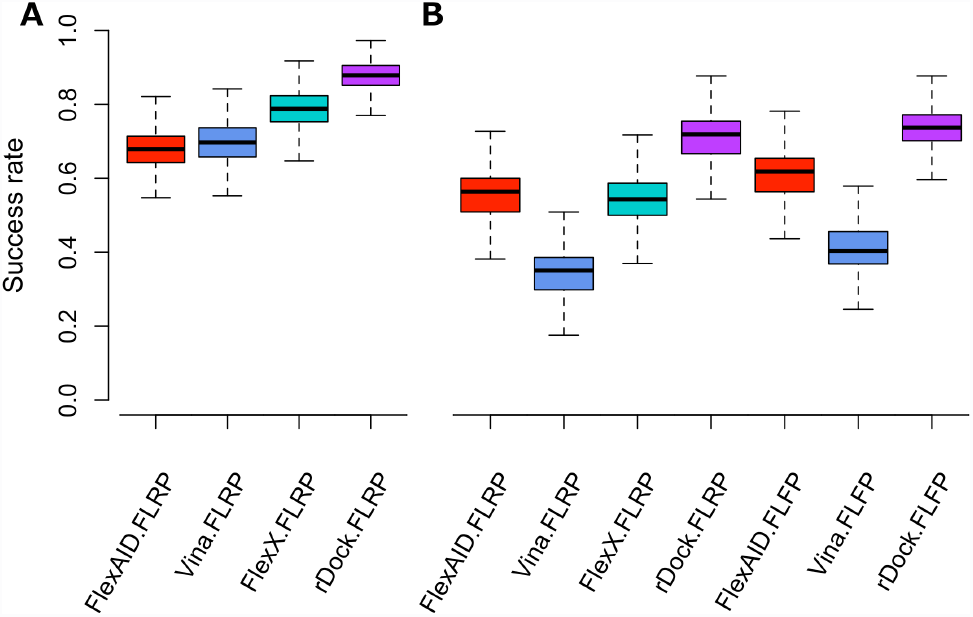
Comparison of the performance of FlexAID, AutoDock Vina, FlexX and rDock for binding mode prediction. The performance of FlexAID (red), AutoDock Vina (blue), FlexX (cyan) and rDock (purple) were tested on the Astex diverse set (A) in the presence of ligand flexibility alone (denoted as FLRP) or presence of ligand and protein flexibility (denoted as FLFP). The programs were also compared on the Astex non-native set (B) in the absence (denoted as FLRP) or presence (denoted as FLFP) of protein flexibility with presence of ligand flexibility. The cases represented are the ideal subset (see Methodology). The results are bootstrapped over 10 000 iterations.

#### Analysis of soft and hard failures in FlexAID

Some of our failures can be attributed to the search procedure, previously categorized as soft failures [36] (not to be confused with the concept of soft scoring functions). We observed that ligands where FlexAID failed to find the correct solution tend to have a larger number of flexible bonds (Figure S5F). For example, FlexAID failed to predict the poses of STI (3-letter ligand residue name) with 7 flexible bonds (PDB ID 1T46), JE2 (PDB ID 1KZK) and BIR (PDB ID 1R1H) with 8 flexible bonds, FR4 with 9 flexible bonds (PDB ID 1UML). The best CF values obtained for these complexes are considerably higher than the CF of the native complex. The simulations were executed with a fixed number of energy evaluations independent of the size of the search space (defined by the volume of the cavity and the number of flexible bonds in the ligand in this case where the target is rigid). These complexes would likely be correctly predicted by increasing the number of energy evaluations.

Other failures are categorized as hard failures as they can be attributed to the scoring. For example, in the case RQ3 (PDB ID 1G9V), the native pose of the ligand is quickly discarded with a considerably higher CF value (*CF*_*ref*_ -89.6) than the predicted pose (*CF*_*min*_ -180.0). In the case of BNE (PDB ID 1MZC), the Zn metal in coordination with the ligand is associated with large steric clashes in the native pose thus lowering the native CF score, a limitation for metals that we wish to address in the future in FlexAID. Another source of missed predictions are cases where the ligand in the native form is solvent exposed. In the case of BSM (PDB ID 2BSM), FlexAID preferentially selects a pose with substantially less hydrophobic surfaces exposed to solvent than observed in the native complex. In some cases such solvent exposure of hydrophobic groups can be offset by other factors. For example, the best predictions for AO5 (PDB ID 1R58) are substantially less exposed to solvent than the native pose (~35Å^2^ vs ~115Å^2^). The native pose in the biological unit exposes to solvent a bulky hydrophobic group found buried in our predictions. In the native complex, the entropic penalty involved in exposing a hydrophobic group to the solvent is partially offset by the enthalpic gains through interactions with 2 Mn ions in the binding-site. Unfortunately, our interaction matrix does not currently appropriately cover interactions with Mn due to a lack of examples of these interactions in the training data (Table S1). Another case of poor parameterization of an uncommon atom is the case of PVB (PDB ID 1V0P). In this case, our failure could be attributed in part to the Cl atom of the ligand. This atom is exposed to the solvent in the native pose and found buried in a hydrophobic cavity in our best prediction, as its exposure to solvent is highly penalized according to the parameter set. Lastly, the solvation term is responsible for a number of other failures. In the case of 5RM (PDB ID 1XM6), FlexAID identifies a pose with nearly identical interactions than the native pose but RMSD of 2.6Å due to the burial of a flexible moiety found more exposed in the native form.

To summarize, most hard failures involve less frequently observed atom types in biological molecules and are caused by the lack of training data given the nature of our approach. As seen, some parameters could be overfitted if the atom types are not sufficiently represented in the decoy sets or could not be optimized due to absence of the atom types in the ligands used. To account for the lack of examples in the training data, we mapped infrequent atom types to a frequent atom type with similar chemical properties or, if none was available, pairwise interactions involving the infrequent atom type were set to zero. The mapping of Zn to Mn allows the complex 1R58 to be properly docked with good ranking (Rank of 3). Predictions are also improved when interactions made with Cl atoms are made to be neutral as in the two cases that follow. In the case of STC (PDB ID 1L2S), a significantly better ranking is observed (rank 1 instead of 51) and for the hard failure PVB (PDB ID 1V0P) discussed above, we successfully predict the native complex as top ranked solution.

Considering the soft nature of our scoring function, one could expect that FlexAID would perform better on predominantly hydrophobic binding-sites where directionality is less important. We do not observe any clear correlation of the rank with the hydrophobicity index of the binding-site (Figures S6A-F).

#### Rigid ligands on flexible non-native structures

We observed above that the AutoDock Vina scoring function failed with rigid ligands when docking into native structures even if target flexibility was not necessary (Figure 4A). At the same time, adding ligand flexibility allows AutoDock to perform well in rigid non-native structures. We performed the opposite experiment, docking rigid ligands on flexible non-native structures to understand if setting the protein as flexible instead of the ligand can have the same effect. It turns out that introducing target flexibility for rigid ligands in their correct conformation does not help recover high success rates with AutoDock Vina (8.8% success rate, Figure S4B). In the same conditions, FlexAID attains a success rate of 83.6% and rDock obtains 78.9% (Figure S4B). We do not include FlexX in this experiment as this program cannot account for receptor flexibility.

#### Flexible ligands on rigid non-native structures

We compared the performance of FlexAID when docking into rigid non-native structures to the ones of AutoDock Vina, FlexX and rDock. For each method, the protein-ligand complexes that could not be successfully predicted in the native structures were not included (see Methodology). Thus, the failures observed in this experiment can be attributed to the conformational changes of the binding-site. When it comes to non native-complex structures the success rate of AutoDock Vina and FlexX decrease considerably to a level below that of FlexAID that is minimally affected by the transition to rigid non native-complex structures (four left-most bars in Figure 4B). FlexAID predicts 56.4% of the complexes when side-chain flexibility is excluded while AutoDock Vina and FlexX and rDock predict 35.1%, 54.3% and 71.9% respectively. A previous study showed that the software GOLD developed by Astex obtains a success rate of 72% on their own Astex non native set [15]. Using the same dataset and criteria for defining success, FlexAID obtains 65.6% when side-chain flexibility is excluded or 71.9% when included. However, we were unable to independently reproduce the predictions reported for GOLD as we were unable to obtain a license.

To further understand the sensitivity of the scoring functions, we calculated the success rate as a function of the magnitude of rearrangements observed between native and non-native forms. When simulating rigid non native-structure targets (Figure 5A), FlexX performs the best in non-native structures that are closely structurally similar to the native form. However, despite the observed decrease in success rate for all programs as the magnitude of movements increase, FlexX has a steeper slope compared to those of FlexAID, AutoDock Vina and rDock. The performance of FlexAID surpasses the one of FlexX on structures with maximal displacement greater than 3Å but remains below that of rDock.

**Figure 5.**
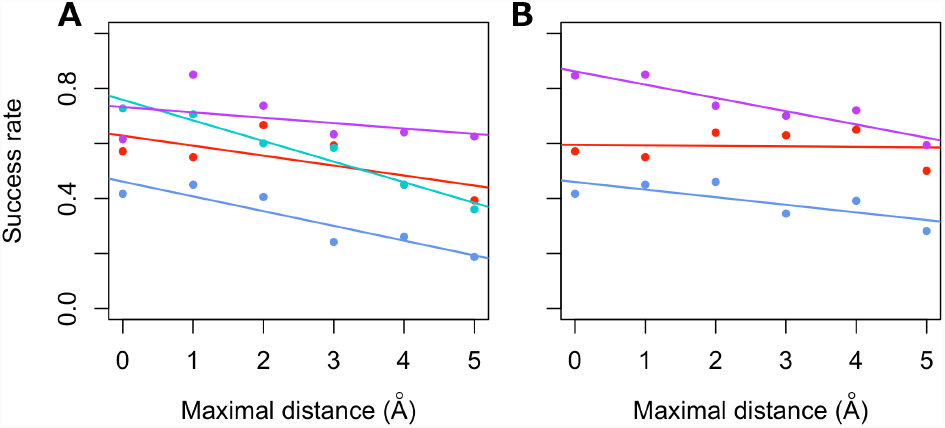

Impact of the magnitude of movements on the success rate. We compare the success rates of FlexAID (red), AutoDock Vina (blue), FlexX (cyan) and rDock (purple) as a function of the maximal displacement of any atom of the binding-site between apo/non-apo and holo forms of the Astex diverse sets. We compare the success rates between the methods in absence (A) or presence (B) of protein flexibility. The following bins were used maximal displacement: [0.0,1.0[, [1.0,2.0[, [2.0,3.0[, [3.0,4.0[, [4.0,5.0[, [5.0,d_max_]. Where d_max_ is the maximum displacement observed across all pairs. Each point is a bootstrapped success rate over 10 000 iterations. The lines represent the linear regression of all points for a given method.

#### Flexible ligands on flexible non-native structures

We compared the performances of FlexAID, AutoDock Vina and rDock when docking into flexible non-native structures. This is the most realistic of all scenarios as it is the situation likely to be found when the structure of a ligand-protein complex is unknown prior to the docking simulation. FlexX considers the protein as rigid, thus we did not include it in experiments involving protein flexibility. A variant of the method called FlexE introduces protein flexibility by combining a set of up to 16 protein structures [14], creating an ensemble that is used to combine structurally variable parts from each structure. The reported results for FlexE using a dataset of 10 examples show the same success rate as that of combining the results of FlexX from all alternative structures docked separately. Therefore, for all purposes, we can consider that the results obtained in the previous section using FlexX on flexible ligands docked against rigid non-native structures reflect the results that would be obtained with FlexE.

The inclusion of side chain flexibility in FlexAID and AutoDock Vina increases their success rates by approximately 5% to 61.8% and 40.4% respectively compared to 73.3% for rDock. However, for maximal displacements above 2Å, rDock and FlexAID achieve consistently comparable success rates (Figure 5B). Interestingly, in this situation the performance of FlexAID is independent of the maximal displacement while other programs are affected by increasing amounts of target variability despite including target flexibility in the simulations. This result suggests that FlexAID may have an advantage as the magnitude of movements required to accommodate the ligand increase. To test this hypothesis, we analysed subsets of the Astex diverse set as a function of the number of side-chains with large unfavourable contributions to the CF function due to steric clashes when superimposing the holo and non-holo forms. We call such side-chains as critical as these must undergo movements to accommodate the ligand. As with previous situations, each program has its own ideal subset of targets. While rDock achieves a 73.3% success rate compared to 61.8% for FlexAID on the entire Astex diverse set, when focusing on cases with at least one critical side-chain, the success rate of rDock decreases to 53.3% (on 494 non-holo targets representing 41 cases) while that of FlexAID is only mildly affected and higher than that of rDock at 58.3% (392 non-holos/36 targets). When looking at three or more critical side-chains, rDock is unable to correctly predict any of the 31 non-holo structures representing 9 targets. Meanwhile, FlexAID has a success rate of 37.5% for 30 non-holo cases representing 8 targets with at least 3 critical side-chain movements. In the past our group developed a non-redundant dataset of apo holo pairs and showed that in 90% of cases at least one side-chain undergoes a rotamer change upon binding and in 30% of cases such movements are critical [6]. It is unclear to what extent either the Astex diverse set or our non-redundant dataset are representative of the true levels of critical side-chain movements but in cases where any number of such movements are present, FlexAID has a clear advantage over rDock.

### Virtual screening

The analysis of a small-molecule docking program cannot be complete without the analysis of its performance on virtual screening. However, it has to be stressed that the prediction of binding poses as performed above and that of discriminating binding from non-binding small molecules are two distinct problems. We evaluated the performance of FlexAID in virtual screening by docking against 38 targets of the DUD dataset [37]. We use the average AUC and enrichment factors at 1% of the ranked database (EF1%) as criteria of performance in virtual screening. In average, we have an AUC of 0.56 (and EF1% of 2.9) across all targets (Figures S7A-B), which closely matches results obtained from a previous study that used DOCK6 [11]. Interestingly, despite the differences in the nature of the two algorithms, some of their best targets (GART, SAHH, RXR and COX2) in terms of AUC are also targets where we had the most success as we obtained AUCs of 0.83, 0.81, 0.92 and 0.80 respectively. Using an AUC of 0.6 as a rough guideline for successful predictions over random, FlexAID and DOCK6 succeed for 16 targets of which 10 are shared. However, our virtual screening results are on average lower than those for rDock, Glide and AutoDock Vina [18] using the complementarity function (CF). These results were somehow expected considering the methodological approach used to derive the CF. As the CF was optimized for scoring appropriately native and near-native binding modes, this may come at the expense of enrichment [30]. We therefore investigated if re-scoring could further enhance the predictions of FlexAID in virtual screening. The re-scoring of docking poses is a common practice to account for the possible caveats and biases of scoring functions [38]. We utilized the free knowledge-based scoring function RankScore as re-scoring method. In average, the AUC and EF1% increased to 0.62 and 4.9 when using RankScore [26] (Figures S7A-B) compared to 0.66 and 8.9 respectively for AutoDock Vina, 0.69 and 11.4 for rDock and 0.78 and 22.6 for Glide [18]. The size of the dataset used to obtain the virtual screening results for AutoDock Vina, rDock and Glide is similar to ours but 20 of the DUD targets were replaced in their case with DUD-E targets. Apart from this potential source of differences, our results are comparable to those of AutoDock Vina and to some extent those of rDock but still considerably lower than Glide.

### Graphical user interface

We developed a graphical user interface (GUI) called NRGsuite that is directly integrated into PyMOL to allow non-experts in the field to easily use FlexAID. Through the GUI, the users can easily set the target and ligand to be docked, define, measure and partition the binding-site volume, view the simulation in real-time, etc. An extensive manual is provided in order to guide the users through the different steps. The NRGsuite is freely available and its installation also installs FlexAID. The NRGsuite is compatible under the operating systems Linux, MacOS X and Windows. A complete description of the NRGsuite features is beyond the scope of the present work and will be fully described elsewhere.

## DISCUSSION

In this paper we present FlexAID, a docking method that uses a soft and smooth scoring function based on surfaces in contact. While all current scoring functions used in major docking algorithms impose strict geometric constraints to define molecular interactions, our scoring function focuses on coarser features such as shape complementary. Our scoring function, together with the unique way in which our pairwise energy terms were derived, contribute to produce better results in binding mode prediction than several widely used methods. As a matter of fact, in the most challenging and realistic of situations, when docking on non native-complex structures, FlexAID outperforms the widely used AutoDock Vina and FlexX (and FlexE by extrapolation) programs and equates the performance of the state-of-the-art rDock program. However, FlexAID outperforms rDock when critical side-chain movements are necessary to accommodate ligand binding. One potential advantage of a program such as FlexAID with a soft, smooth scoring function over programs that utilize hard scoring functions is that the former are less sensitive to structural variations such as those that are common in homology models. A well-characterized phenomenon in docking programs with hard scoring functions is that the accuracy of the results is on average higher for holo form targets, followed by non-apo form targets and lastly homology models [39]. While it remains to be tested, given the soft and smooth scoring function in FlexAID, the program may be well suited for binding mode predictions and virtual screening using homology models.

Despite that more iterations were carried out to improve our scoring function (data not shown), we did not observe any further improvements on the training set in terms of success rate and ranking. We may have achieved a limit for the optimization of the scoring function with the fixed set of atom types used. However, it may be possible to augment the number of atom types. In fact, inaccuracies in the Sybyl atom types have previously been discussed [40]. For instance, the O.3 atom type definition could be divided in at least 2 atom types as it accounts for both hydroxyl and ether groups. Such a separation is chemically meaningful as only hydroxyls can act as H-bond donors. The knowledge-based scoring function DrugScoreX increased its atom types library size to over 150 from its original version. Our scoring function would most likely be improved by extending the number of atom types as that would allow us to define more appropriately the chemical properties of atoms. Moreover, most of the hard failures observed are due to the absence or lack of sufficient training data (Table S1). Therefore, whereas in some cases it is chemically plausible to split an atom type into two, in other cases, it may be worth merging atom types with similar chemical properties due to the lack of sufficient data.

The interaction between atom types was derived from an iterative procedure using both positive and negative information with large decoy sets. In the present study we utilize poses of a ligand with RMSD values higher than 2Å compared to its experimentally determined pose to define negative decoys. However, the combination of decoys for different ligands as negative decoys for each target may further improve virtual screening [41]. It is essential to couple the sampling and scoring steps, i.e. the decoys introduced in the decoy sets must be representative of poses that are likely to be scored during the docking simulations [42]. Despite the fact that none of the decoys included protein flexibility, our scoring function could still maintain good performance when docking into non-native proteins. Future decoys sets will include side-chain and backbone flexibility. Thus, one major advantage of a soft scoring function over a hard one is that it can implicitly account for protein flexibility. This smoothness cannot be explained alone by the softening of the steric clash term [43] but is also acquired through the use of surfaces in contact as the smoothness of the CF is gradually increased through the iterative process. The values of smooth scoring functions do not drastically change with small rearrangements of the structure. Therefore, achieving the global minimum of the CF in a traditional genetic algorithms docking simulation can be done within less energy evaluations than with hard scoring functions. All simulations with FlexAID were performed with 10^6^ energy evaluations while at least 25 times more energy evaluations were used in the case AutoDock Vina [12]. Despite that more time is required in computing surfaces in contact than simple distance calculations, the extra time required is offset by the smaller amount of energy evaluations required. Therefore, there appears to be a trade-off between the search and the scoring that can quickly become computationally intensive as more and more degrees of freedom are required to account for the stringent geometric constraints. Because the search is more complex when hard scoring functions are used, traditional genetic algorithms have included additional minimization steps in order to fulfill these constraints [44], something that is not required with FlexAID.

Given that the best RMSD obtained correlates with the number of rotatable bonds in the ligand, improvements in the methodology used to simulate ligand flexibility may increase the accuracy of FlexAID. Currently each flexible ligand bond represents a genetic algorithm variable sampling angles at 10° intervals and intramolecular ligand interactions do not contribute to the scoring. A number of different possibilities may be explored in the future to address the gap in the loss of success rate such as finer sampling, inclusion of intramolecular ligand interactions in the CF function as well as the possibility of pre-computing favourable ligand conformations [45, 46]. Users of FlexAID have full control over ligand flexibility and as such may choose in addition to allowing it in FlexAID, also docking pre-generated ensembles of ligand conformations as rigid ligands on flexible non-native targets as in this case we obtain 83.6% success rate compared to 78.9% for rDock.

In terms of virtual screening, FlexAID requires rescoring to achieve comparable results to those of AutoDock Vina and rDock (although different datasets were used as noted, and a direct comparison may not be straightforward). Considering the differences between binding pose prediction and virtual screening and our emphasis in the development of the scoring function for the former, the result obtained are expected but may be further improved with changes in the scoring function discussed above. Lastly, it is important to keep in mind that the above experiments were performed with flexible ligands and rigid targets and improvements may be observed with the inclusion of target flexibility as it was previously observed that multiple receptor conformations lead to improvements [47].

In this work we introduce FlexAID, a small-molecule docking algorithm with a smooth soft scoring function. We compare FlexAID to commonly used (AutoDock Vina, FlexX) and state-of-the-art (rDock) software in the field as well as to reported results for Glide and GOLD and DOCK6. The comparisons on the prediction of binding poses are carried out using the accepted *de facto* benchmarking Astex diverse set, which offers an independent dataset (except for GOLD developed by Astex) to test docking programs. Notably missing are a number of commercially available software that are widely used such as GOLD and Glide among others for which we could only minimally compare to published data. It would be useful if commercial software developers would at least make all data available for such benchmark datasets to allow more detailed comparisons even if independent validation is not possible.

Our results show that FlexAID is a highly competitive mature docking program that performs better than several widely used methods and uniquely apt at docking ligands in flexible binding-sites where flexibility is crucial for binding. FlexAID high-performance and availability in all platforms, particularly Windows will considerably increase the reach of docking simulations as a research and teaching tool.

## METHODS

### Binding-site definition

There are two methods to define the spatial volume that a ligand can occupy during search in FlexAID. The first and most simplistic is the definition of a sphere with radius and center coordinates defined by the user. This method is not efficient as invariably a large volume of the search sphere is in practice inaccessible to the ligand due to steric clashes with protein atoms. The second method involves the detection and use of one or more cavities of the target molecule. A grid with spacing of 0.375Å is built within the volume of each cavity to define the searchable area of the ligand where each grid vertex serves as an anchor point for the reference atom of the ligand. This discretization of space allows us to replace three translational degrees of freedom for a single variable. When the ligand is processed, the reference atom is internally defined and is normally identified as an atom within the largest rigid portion of the ligand. The ligand will not necessarily be fully contained within the volume of the binding-site, as only the reference atom is required to be anchored to a grid point. This permits us to account for remodelling of the binding-site such as when side-chain flexibility is included while maintaining the originally defined grid.

The use of cavities is preferred as it permits an accurate definition of binding-sites reflecting precisely the complex geometry of each cavity. Cavities are calculated using our own implementation of the SURFNET algorithm [48] called GetCleft. Briefly, the method inserts spheres between each pair of atoms of the target and reduces their radii until there are no more clashes. A cleft is defined as an ensemble of connected spheres. This binding-site definition is similar to the definition in the program DOCK [49]. One or more cavities can be searched at the same time. The possibility to perform a global search against all cavities may be interesting when searching for binding hotspots [50] or druggable allosteric binding sites.

### The scoring function

We use a modified form of the complementarity function to evaluate the energy of the complex [21]. A subset of 40 Sybyl atom types [51] is used to describe the chemical properties of atoms. We assign atom types using the open-source and freely accessible software Open Babel [52]. The Van der Waals (VDW) radii of heavy atoms are expanded to implicitly include Hydrogen atoms [53]. The VDW radius of each atom is further expanded by the radius of a water molecule (1.4Å) to calculate contact surfaces between atoms and solvent accessible surfaces (SAS) with the implicit solvent. We calculate surface areas in contact analytically using the constrained Voronoi procedure of McConkey *et al.* [54]. The scoring function is given by:

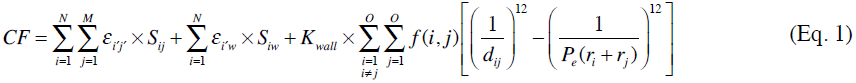

where atoms *i* and *j*, with radii *r*_*i*_ and *r*_*j*_ belong to the ligand and protein with *N* and *M* atoms respectively. The interaction energy between atoms *i* and *j* of Sybyl atom types *i′* and *j′*, is given by *ε*_*i′j′*_ and modulated by their surface area in contact *S*_*ij*_. Similarly to GOLD and DOCK, intramolecular interactions are omitted. The second term is used to simulate the hydrophobic effect, through an effective interaction energy between ligand atoms and the solvent *ε*_*i′w*_ modulated by the solvent accessible area *S*_*iw*_ of each ligand atom. As this term involves only ligand atoms, we assume that different poses of the ligand desolvate the target equally [55]. The last term accounts for the repulsive VDW interactions. The constant *K*_*wall*_ = 10^6^, is used to penalize steric clashes when the distance between atoms *d*_*ij*_ is smaller than the sum of their van der Wall radii. The permeability factor, *P*_*e*_ = 0.9, softens the potential to allow some receptor plasticity [43]. The repulsive VDW interactions are only calculated between non-covalently bonded atoms and atoms separated by at least 3 consecutive covalent bonds similarly to other methods [56]. The function maximizes the area in contact between atoms with favourable interaction energies while minimizing solvent exposed areas and steric clashes.

The *ε*_*i′j′*_ interaction energies are such that negative values represent favourable interactions and were obtained using an iterative optimization approach. Monte Carlo simulations were used to optimize the parameters such that they can discriminate, in as much as possible, between native and near-native binding modes (referred as true positives) from low-energy decoys (referred as false positives). During the Monte Carlo optimization, the probability of changing the value of a pairwise parameter was proportional to the frequency that it was observed in the PDBbind dataset used in training to guide the search more efficiently. We used the Area Under the Curve (AUC) of Receiver Operator Curves (ROC) as the objective optimization function. Independent datasets were used in the Monte Carlo optimizations and in the validation the resulting potentials. We iteratively used the best set of interactions to enrich the decoys set with harder false positive decoys (lower CF values) as well as increased the number of complexes (and their respective decoy sets) in each consecutive Monte Carlo optimization (Table 1).

Decoy sets were generated using FlexAID for each iteration with the best potential obtained in the previous iteration. We utilized the PDBbind [27] refined-set (release 2012) to predict the native binding modes of an ensemble of crystal structures representing protein-ligand complexes. The proteins were kept rigid but ligands were fully flexible. We excluded complexes for which our method could not successfully predict the native binding mode. The remaining complexes (Table 1) had at least 2 true positives (the native pose of the ligand from the crystal and a near-native binding mode) and an unlimited number of false positives. Each complex utilized had it own decoy set representing different conformations and relative positions of the same ligand against its own receptor. We used thresholds of RMSD ≤ 1.5Å for ligands with less than 15 heavy atoms [57] or RMSD ≤ 2.0Å otherwise to define true positive decoys. Every decoy had to have an RMSD of at least 0.5Å from any other decoy in its decoy set. Complexes in the PDBbind dataset that also appear in the Astex diverse or non-native sets were removed. Lastly, in order to prevent a bias towards interactions that are highly present in particular protein families, we grouped proteins by sequence identity to minimize redundancy. Each protein group was given an equal weight during the optimization. We aligned protein sequences using ClustalW [58]. Proteins belonged to the same group if they shared at least 30% sequence identity. The final set of pairwise energy terms used in FlexAID is presented in Table S3.

### General features of FlexAID

FlexAID is a probabilistic docking program that optimizes the ligand-protein complex by minimizing the complementarity function using genetic-algorithms (GA) as search procedure. A ligand-protein conformation is referred as an individual within a population. Each individual is constituted of one chromosome. Each gene represents one optimization variable. Four genes account for the rotation and translation of the ligand. Each flexible ligand dihedral bond represents an extra gene. Briefly, the GA optimization works in 6 steps. 1. An initial population of individuals is generated randomly. 2. The complementarity function (CF) evaluates the energy of the whole population. 3. The individuals are ranked according to a fitness function. 4. The fittest individuals, having a greater probability to reproduce, are selected pairwise to produce two new offspring. 5. A new population is created according to a reproduction technique. 6. Loop through steps 2 to 5 with the new population. The population converges towards a minimum as the generations increase. We implemented an adaptive GA to maintain diversity in the population and prevent early converge to local minima [59]. The probability of the genetic operators (mutation and crossover) increase when the population converges to a (potentially local) minimum to generate additional conformational diversity in the population and escape the minimum found. Given the properties of the selection technique (see below), a minimum will remain in the population until a lower minimum is found or the maximum number of generations is run.

The user can choose between linear or shared (default) fitness functions. Fitness functions are mappings that permit to weight individuals according to their CF values to further generate diversity in the population and prevent premature converge. For linear fitness, the fitness grows linearly as the CF decreases and acts as a scaling factor. The shared fitness function mimics nature, where individuals sharing a specific geographical niche (in our case similar pose) need to share resources. Internally, the fitness of individuals sharing similar search space is lowered. Two different reproduction techniques are available to the user: Steady State and Population Boom (default). While the first one is fairly well documented, the latter is a reproduction technique first introduced in the original version of FlexAID [29]. Briefly, Population Boom creates an entirely new population with N individuals according to the standard rules to create offspring then combines the old and new populations and selects the best fit N individuals out of the 2N. If we denote the fitness of an individual *i* in generation *t* as 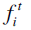; whereas standard reproduction techniques ensure 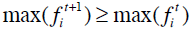 Population Boom ensures by definition that 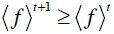. In plain words, instead of ensuring that a small number of the best-fit individual(s) are kept between two consecutive GA generations, Population Boom ensures that the average fitness of the whole population will not decrease through the generations.

FlexAID takes as input a configuration file and a GA parameters file. The GA file allows the user to adjust the population size and the number of generations to control the length of simulations, to control the parameters of the genetic operators to make them adaptive (default) or constant and to change the reproduction technique and the fitness function. The configuration file is highly customizable and is used to specify the target and the ligand to be docked, define the binding-site, include extra degrees of freedom representing ligand and protein flexibility and modify internal parameters of the program. The target and ligand files need to be processed with our auxiliary program Process_Ligand in order to derive the Sybyl atom types. PDB, MDL and MOL2 formats are supported for the definition of the ligand and target. The processed ligand file encodes an internal coordinates system used to incrementally build the ligand into the binding-site. FlexAID generates a fixed but customizable number of results written in PDB format (default is 10). These represent a geometric clustering of the individuals sampled during the whole simulation. For example, in the hypothetical case where the entire population converges to a single solution at the end, other sampled good solutions that didn’t necessarily survive to the last generation are brought back to life and added to the list of the top ten solutions according to their scoring. The user has the option to only output atoms that could move during the simulation or output the whole complex. The latter is essential if the user desires to re-score complexes with external scoring functions when side-chain flexibility is allowed, a missing feature even in modern docking suites [18]. One final characteristic of FlexAID is that in addition to proteins, also nucleic acids (both RNA and DNA) can be used as rigid targets.

### Side-chain flexibility

FlexAID uses a flexibility probability scale for side-chain rotamer changes [6] (Table S4). The probability scale serves as a scaling factor to the mutation operator. Whilst the probability of undergoing conformational changes is used to bias the sampling of flexible residues, the user still needs to explicitly define the list of flexible side-chains. Our analysis [6] offers a number of pointers to choose the nature and number of flexible residues. Users should preferentially choose side-chains that are highly exposed to solvent and when working with X-ray structures, that have high b-factors. Moreover, we observed that a maximum of 5 flexible side-chains account for approximately 90% of binding-sites. Side-chains are modeled using rotamers defined in the Penultimate Rotamer Library [60]. For any given amino acid, all side-chain rotamers are considered equally probable despite the distinct probabilities of rotamers. Each flexible side-chain represents one extra gene following an indexing of rotamers.

### Docking experiments

We evaluate our method for the prediction of binding modes using the Astex diverse set comprising 85 protein-ligand complexes as well as the Astex non-native set comprising 1112 structures representing apo and non-holo forms of 65 proteins from the diverse set [15, 31]. The complex with PDB code 1OF6 was removed due to errors in defining atom types. We include side-chain flexibility when docking into the non-native forms of the proteins. AutoDock Vina and FlexAID describe each degree of freedom individually. For FlexAID and AutoDock Vina, the set of flexible side-chains in this experiment was restricted to those that are required to move to sterically accommodate ligand binding. Such critical side-chains are those where large steric repulsion was observed when the ligand is superimposed into the apo or non-holo forms as previously defined [6]. While the explicit selection of side-chains represents a bias on its own, it allows us to do a controlled experiment limiting the search space in a focused manner. Lastly, in rDock protein flexibility is included in the form of rotations of polar Hydrogen atoms of binding-site side-chains.

Considering that the native pose of ligands is known in the set of complexes used to validate the prediction of binding mode, the search space is reduced by restraining the volume of the binding-site to a given volume surrounding the ligand in the native pose. In the docking experiments described in this work, clefts defined with GetCleft were restricted to retain grid vertices within 5Å of any atom of the ligand. We use the same binding-site definition when docking into the non-native forms as well as for the virtual screening experiments. The binding-site definition of FlexX is equivalent as it is defined as the set of protein atoms contained within 5Å of any atom of the ligand. We define the binding-site for rDock using the cavity in which the reference ligand is contained. AutoDock Vina defines a binding-site using a rectangular box [12]. In order to use a binding-site definition equivalent to that of AutoDock Vina, we defined the rectangular box as the smallest one that enclosed the cleft used in the FlexAID simulations. Although this binding-site definition may appear as a fair comparison in terms of search space, we obtain poor results with AutoDock Vina with this definition of binding-site. When expanding the rectangular box by 10Å in each dimension we obtain significantly better performance on both native and non native forms, comparable to results obtained by others using AutoDock Vina [18]. We increased the exhaustiveness to 16 compared to the default value of 8 when docking with AutoDock Vina as previously done [18]. We use the threshold RMSD ≤ 2.0Å as measure of success in binding mode prediction.

Success rates were bootstrapped [35] over 10000 iterations. A program may fail on a particular target for technical reasons. Therefore, we bootstrapped the results for each program individually with its own subset of the Astex dataset to provide the best possible estimation of the success rate for each method. In the results section, we also refer to the largest common subset, i.e. the subset comprising only cases that were docked with all programs. In the non-native docking experiments, for each method we remove from its own subset, proteins that could not be predicted in the native form. This allows us to exclude cases that may fail because of variables other than protein flexibility. We apply the latter filter as we feel that the purpose of non-native docking experiments is to assess the influence of protein flexibility. The Astex non-native set is biased as it contains proteins that were crystallized more often than others. To account for this bias, the bootstrapping procedure is done in 2 steps with the selection of proteins followed by the selection single non-native conformers for the given protein thus not biasing the estimation of success rates with over-represented cases given the combinatorial combination structures.

We implemented a grid-computing framework using the BOINC infrastructure [61]. The project NRG@Home (for Najmanovich Research Group at Home) allows people around the world to contribute to our work by allowing their computers to perform FlexAID simulations using the BOINC screen saver. More information on the project and how to join is available at http://bcb.med.usherbrooke.ca/boinc.php. All docking experiments described in this work were run on NRG@Home with a fixed number of generations (1,000) and chromosomes (1,000) for a total of 1,000,000 energy evaluations. Due to the probabilistic nature of genetic algorithms as well as to ensure the return of results from BOINC users, we repeat each docking experiment for a given protein-ligand complex 10 times.

### Availability

FlexAID and its accessory software (GetCleft and Process_Ligand) as well as the NRGsuite are free and open-source and are licensed under the GNU General Public License 3.0. All software is compatible under Windows, MacOS X and Linux. The sources and pre-compiled bundles are available for download at http://bcb.med.usherbrooke.ca/FlexAID where newer version may be found or for the current version also as supplementary information. Information is also provided as supplementary information describing all the necessary steps to execute FlexAID in a command-line manner as well as within the NRGsuite.

## ACKNOWLEDGEMENTS

We are very grateful to all the BOINC users who contributed their CPU time to this work. RJN is part of CR-CHUS, a member of the Institute of Pharmacology of Sherbrooke, PROTEO (the Québec network for research on protein function, structure and engineering) and GRASP (Groupe de Recherche Axé sur la Structure des Protéines).

---

**Table S1.** Atom types and their frequency in the training and test sets

| **SYBYL**<br>**Atom**<br>**type** | **C.1** | **C.2** | **C.3** | **C.ar** | **C.cat** | **N.1** | **N.2** | **N.3** | **N.4** | **N.ar** |
| --- | --- | --- | --- | --- | --- | --- | --- | --- | --- | --- |
| PDBbind | 56 | 10,933 | 44,032 | 28,904 | 875 | 23 | 674 | 0 | 1,478 | 3,096 |
| Astex | 4 | 604 | 2,767 | 1,877 | 36 | 2 | 14 | 18 | 60 | 225 |
|  | **N.am** | **N.pl3** | **O.2** | **O.3** | **O.co2** | **O.ar** | **S.2** | **S.3** | **S.o** | **S.o2** |
| PDBbind | 10,288 | 3,616 | 11,411 | 7,308 | 6,562 | 0 | 17 | 1,208 | 266 | 0 |
| Astex | 569 | 193 | 736 | 330 | 285 | 0 | 6 | 96 | 11 | 1 |
|  | **S.ar** | **P.3** | **F** | **Cl** | **Br** | **I** | **Se** | **Mg** | **Sr** | **Cu** |
| PDBbind | 0 | 367 | 322 | 185 | 63 | 13 | 1 | 67 | 1 | 4 |
| Astex | 0 | 6 | 25 | 17 | 1 | 0 | 0 | 3 | 0 | 0 |
|  | **Mn** | **Hg** | **Cd** | **Ni** | **Zn** | **Ca** | **Fe** | **Co.oh** | **Du** |  |
| PDBbind | 18 | 2 | 1 | 2 | 210 | 30 | 0 | 3 | 0 |  |
| Astex | 2 | 0 | 0 | 0 | 14 | 1 | 3 | 0 | 0 |  |

**Table S2.**
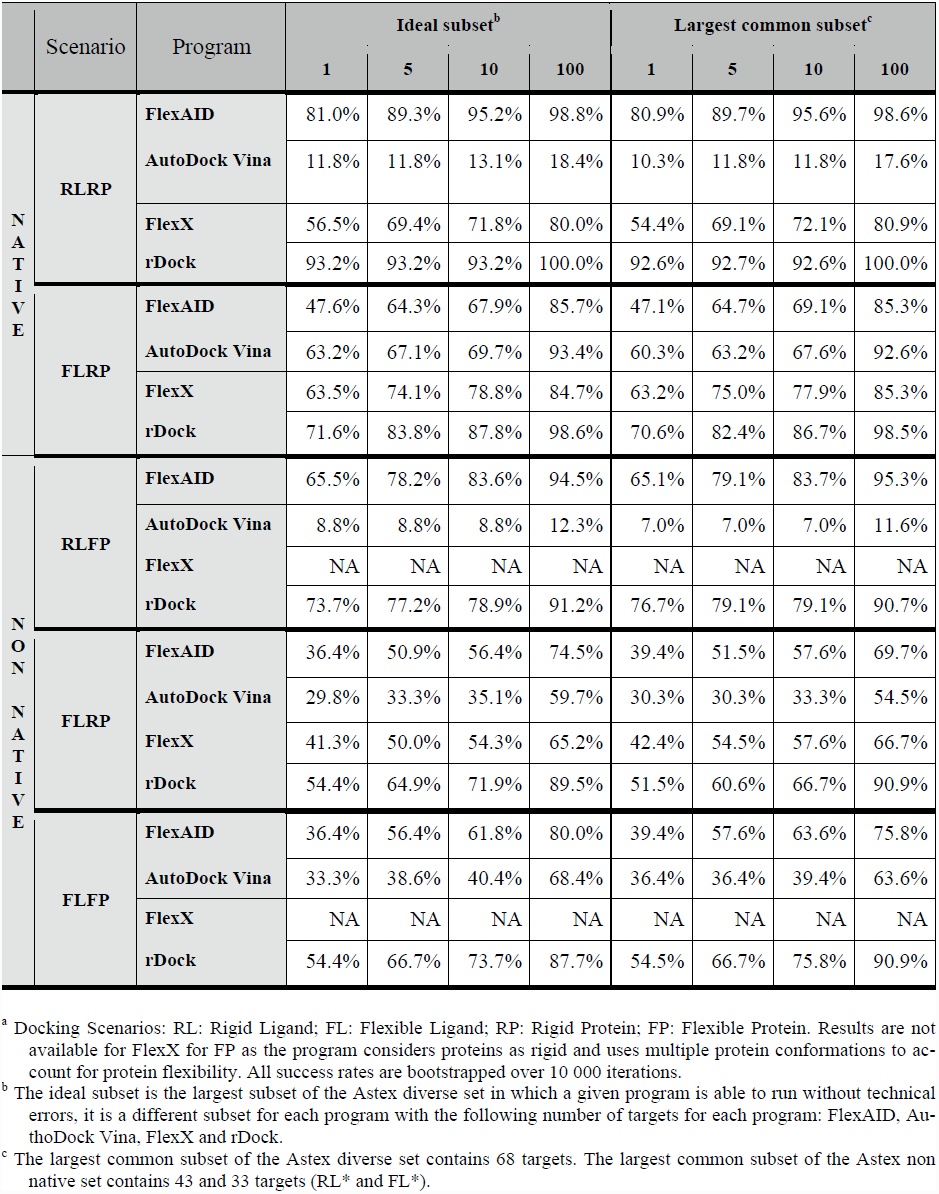
Success rates on top 1, 5, 10 and 100 solutions on different docking scenariosa

**Table S4.** Internal probabilities of rotamer changes for protein side-chains

| **Residue<sup>a</sup>** | **Probability** |
| --- | --- |
| ARG | 0.381 |
| ASN | 0.279 |
| ASP | 0.220 |
| CYS | 0.048 |
| GLN | 0.347 |
| GLU | 0.188 |
| HIS | 0.168 |
| ILE | 0.172 |
| LEU | 0.136 |
| LYS | 0.532 |
| MET | 0.342 |
| PHE | 0.203 |
| SER | 0.195 |
| THR | 0.058 |
| TRP | 0.090 |
| TYR | 0.190 |
| VAL | 0.070 |

<sup>a</sup> Gly and Ala are excluded as their side-chain has no flexible bond as well as Pro as movements in the side-chain inevitably accompanies backbone movements.

**Figure S1.**
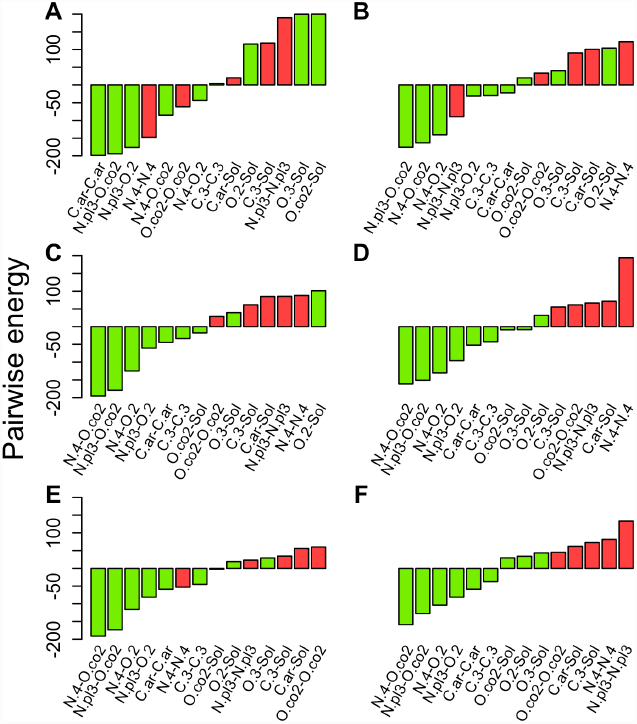
Pairwise interaction parameters for selected interactions for the potentials derived in consecutive MC iterations. Normalized values of the initial potentials (A) and potentials at iterations 1 (B), 2 (C), 3 (D), 4 (E) and 5 (F) are displayed. For the colour coding and atom types nomenclature, refer to the legend of Figure 2.

**Figure S2.**
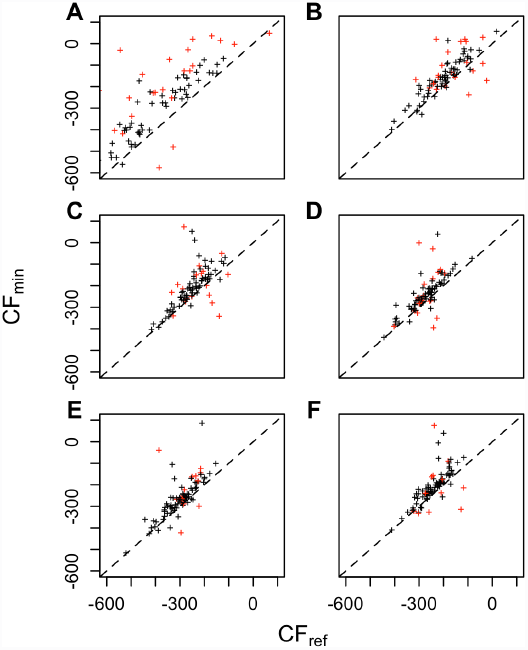

Best CF (*CF*_*min*_) predicted as a function of the reference CF (*CF*_*ref*_). Values for the initial potentials (A) and potentials at iterations 1 (B), 2 (C), 3 (D), 4 (E) and 5 (F) are displayed. Each point represents a unique protein-ligand complex in the Astex diverse set. Points above the dashed line are cases that were optimized relative to the reference CF. Points marked in red are complexes for which no success was observed.

**Figure S3.**
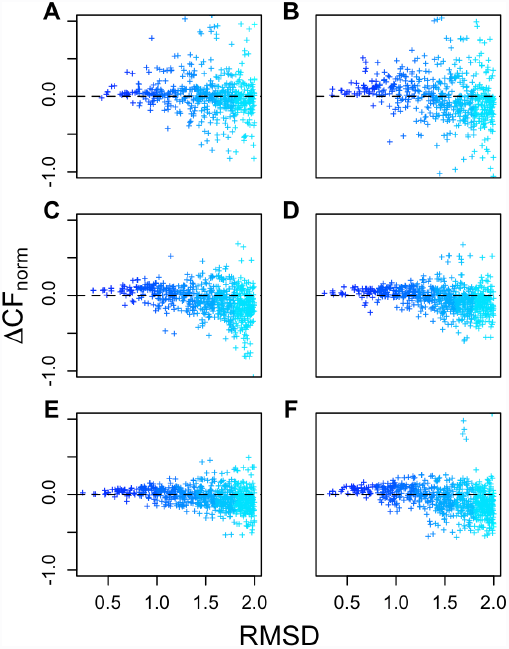

Smoothness of the scoring function for each iteration. Differences in CF between successful poses relative to the reference pose for the initial potentials (A) and potentials at iterations 1 (B), 2 (C), 3 (D), 4 (E) and 5 (F) are displayed. Each point represents a successful prediction in the Astex diverse set. For the colour coding and further detail refer to the legend of Figure 3.

**Figure S4.**
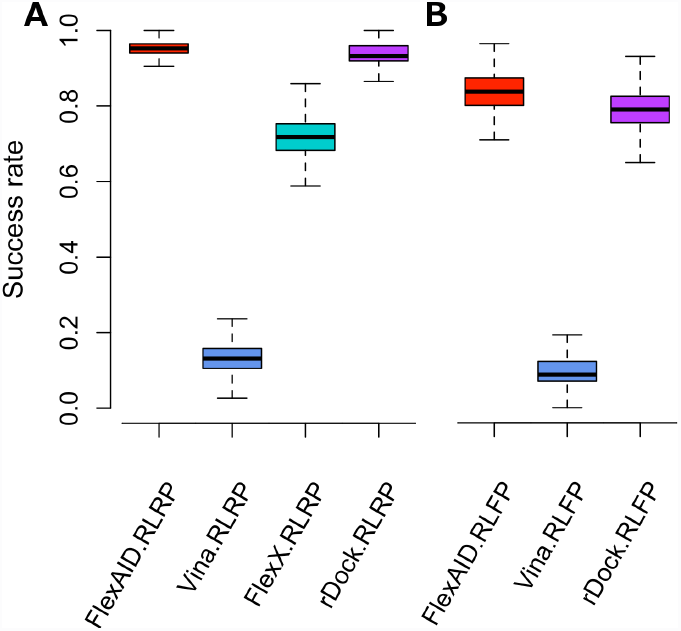
Comparison of the performance of FlexAID and AutoDock Vina for binding mode prediction. The performance of FlexAID (red) and AutoDock Vina (blue) were tested on the Astex diverse set (A) and Astex non native set (B) in the absence of ligand flexibility and receptor flexibility (denoted as RLRP). The cases represented are the ideal subset (see Methodology). The results are bootstrapped over 10 000 iterations.

**Figure S5.**
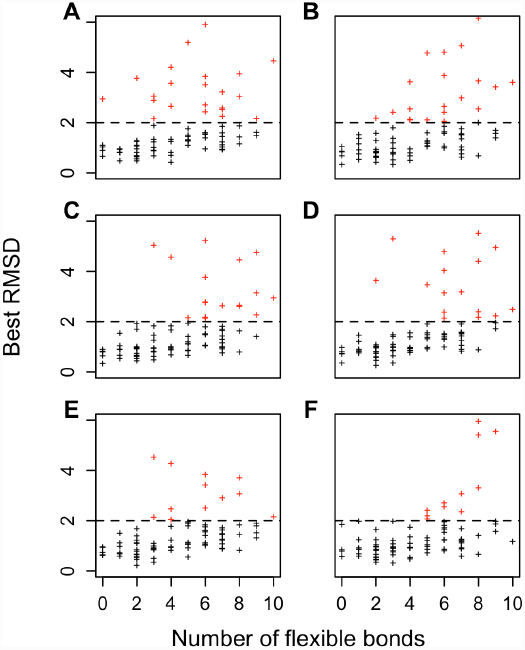
Best RMSD as a function of the number of flexible bonds of the ligand. Values for the initial potentials (A) and potentials at iterations 1 (B), 2 (C), 3 (D), 4 (E) and 5 (F) are displayed. Each point represents the best-predicted pose in terms of RMSD for each complex in the Astex diverse set. A RMSD of 2.0Å is used as cut-off for successful predictions. Points marked in red are failures.

**Figure S6.**
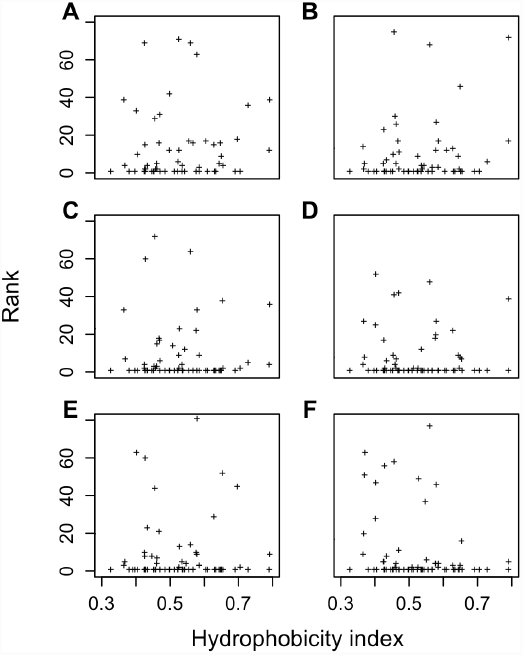

Rank as a function of the hydrophobicity index of the binding-site. Values for the initial potentials (A) and potentials at iterations 1 (B), 2 (C), 3 (D), 4 (E) and 5 (F) are displayed. The hydrophobicity index of a binding-site is defined as the hydrophobic SAS over the total SAS. Each point represents a unique protein-ligand complex in the Astex diverse set.

**Figure S7.**
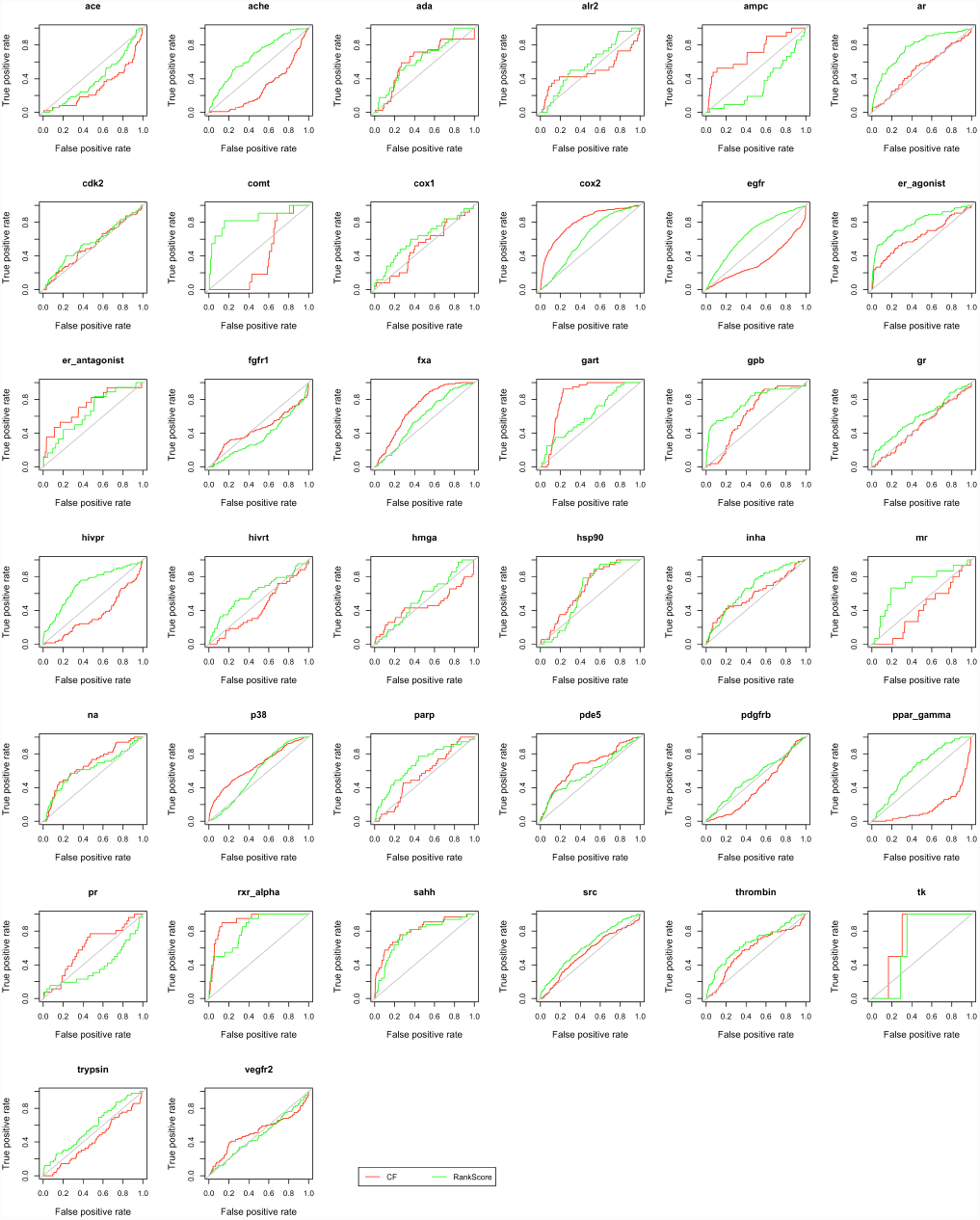
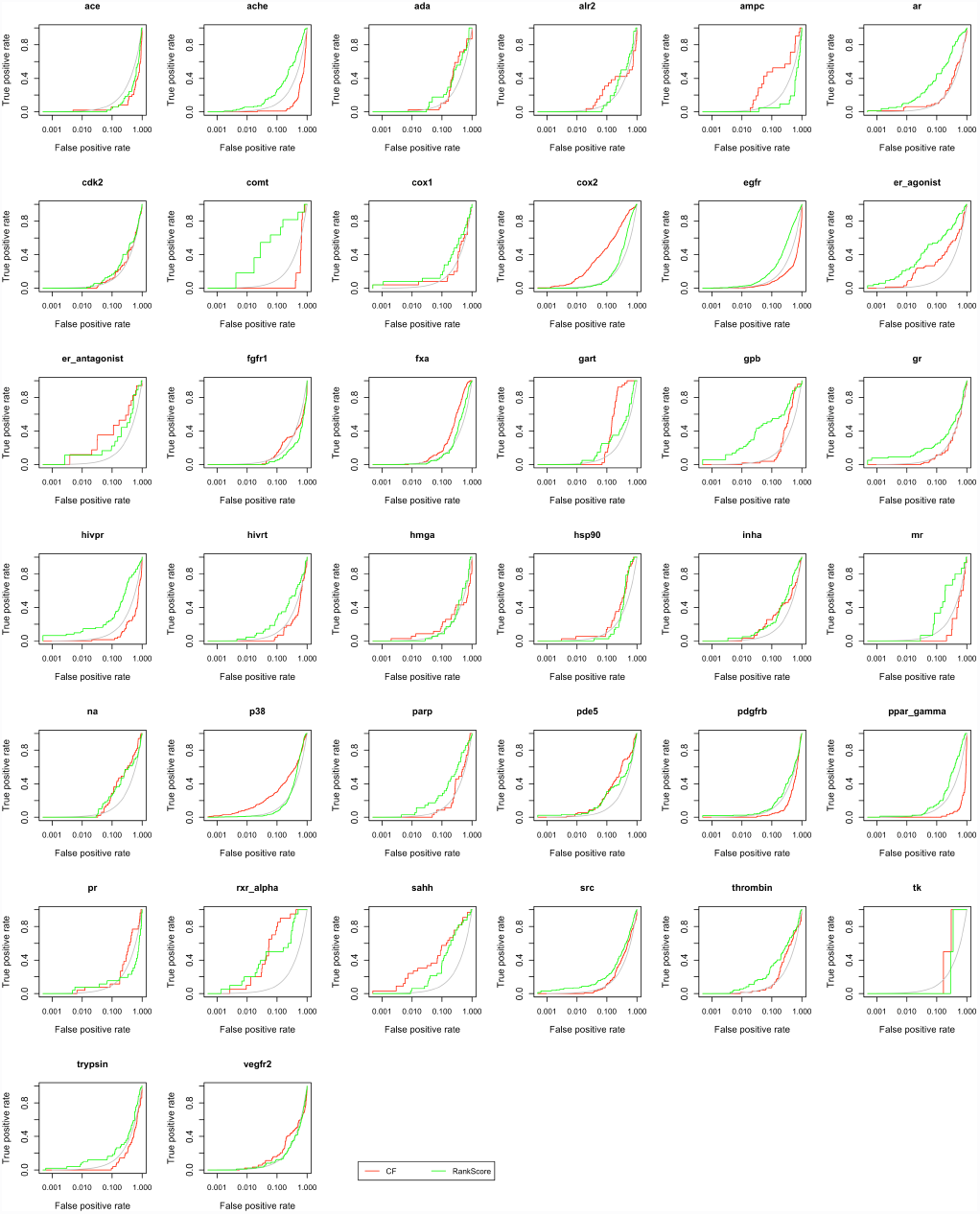

ROC curves of targets in the DUD dataset. The ROC (A) and semi-logarithmic ROC (B) curves of 38 targets in the DUD dataset are displayed. We use the CF (red) and RankScore (green) as scoring functions to discriminate true from false positives. We use RankScore only to re-score the best predicted pose for each unique molecule according to the CF. The grey bars indicate random predictions.

